# Superantigens promote *Staphylococcus aureus* bloodstream infection by eliciting pathogenic interferon-gamma (IFNγ) production that subverts macrophage function

**DOI:** 10.1101/2021.08.17.456537

**Authors:** Stephen W. Tuffs, Mariya I. Goncheva, Stacey X. Xu, Heather C. Craig, Katherine J. Kasper, Joshua Choi, Ronald S. Flannagan, Steven M. Kerfoot, David E. Heinrichs, John K. McCormick

## Abstract

*Staphylococcus aureus* is a foremost bacterial pathogen responsible for a vast array of human diseases. Staphylococcal superantigens (SAgs) constitute a family of potent exotoxins secreted by *S. aureus*, and SAg genes are found ubiquitously in human isolates. SAgs bind directly to MHC class II molecules and T cell receptors, driving extensive T cell activation and cytokine release. Although these toxins have been implicated in serious disease including toxic shock syndrome, we aimed to further elucidate the mechanisms by which SAgs contribute to staphylococcal pathogenesis during septic bloodstream infections. As most conventional mouse strains respond poorly to staphylococcal SAgs, we utilized transgenic mice encoding humanized MHC class II molecules (HLA-DR4) as these animals are much more susceptible to SAg activity. Herein, we demonstrate that SAgs contribute to the severity of *S. aureus* bacteremia by increasing bacterial burden, most notably in the liver. We established that *S. aureus* bloodstream infection severity is mediated by CD4+ T cells and interferon-gamma (IFNγ) is produced to very high levels during infection in a SAg-dependent manner. Bacterial burden and disease severity were reduced by antibody blocking of IFNγ, phenocopying isogenic SAg deletion mutant strains. Additionally, cytokine analysis demonstrated that the immune system was skewed towards a proinflammatory response that was reduced by IFNγ blocking. Infection kinetics and flow cytometry analyses suggested this was a macrophage driven mechanism, which was confirmed through macrophage depletion experiments. Further validation with human leukocytes indicated that excessive IFNγ allowed *S. aureus* to replicate at a higher rate within macrophages. Together, this suggests that SAgs promote *S. aureus* survival by manipulating immune responses that would otherwise be effective at clearing *S. aureus*. This work implicates SAg toxins as critical targets for preventing persistent or severe *S. aureus* disease.

## Introduction

*Staphylococcus aureus* is an important bacterial pathogen that primarily exists as a harmless commensal. Yet, once primary barriers have been breached, this pathobiont also has the propensity to cause an extraordinary range of superficial, invasive, and toxin-mediated diseases (Tong et al., 2015). This spectrum of disease can range from relatively simple soft tissue infection, to pneumonia or bacteremia that may lead to life-threatening sepsis (Kwiecinski and Horswill, 2020; Tong et al., 2015). *S. aureus* is one of the most common causes of sepsis and carries a high mortality rate of 20-40% and mortality rates can double in the context of septic shock (Corl et al., 2020; Kwiecinski and Horswill, 2020). Wide-spread drug resistance, including both hospital and community-associated methicillin-resistant *S. aureus* (MRSA), has further exacerbated treatment challenges with this important pathogen (Turner et al., 2019).

Key to the success of *S. aureus* as a pathogen is a vast array of virulence factors encoded both within the chromosome and on mobile genetic elements. These factors fall into several functional classes including: adhesion factors (e.g. fibronectin-binding proteins A and B [FnbpA and FnbpB]) (Foster et al., 2014; Josse et al., 2017), immunomodulatory proteins (e.g. Staphylococcal protein A [Spa] or Chemotaxis inhibitor protein of *Staphylococcus* [CHIPS]) (Koymans et al., 2017), cytolytic toxins (e.g. Alpha-hemolysin [Hla] (Alonzo and Torres, 2014; Berube and Wardenburg, 2013)) and superantigens (SAg). The SAg family in *S. aureus* consists of at least 26 different paralogues that function by cross-linking major histocompatibility complex (MHC) class II molecules with the variable region of the T cell receptor (TCR) β-chain (Vβ); the unconventional interaction with these two key immune receptors occur irrespective of the cognate antigen and results in the aberrant and widespread activation of T cells followed by proinflammatory cytokine release (Tuffs et al., 2018).

SAgs are the etiological agent of toxic shock syndrome where T cell activation caused by SAgs released from *S. aureus* triggers a systemic ‘cytokine storm’ that can lead to hypotension and multiple organ failure, and in some cases death. SAg activity has also been implicated in a number of other serious diseases including endocarditis, pneumonia and bacteremia (Spaulding et al., 2013). Historically, it has been difficult to model the biological functions of SAg activity *in vivo* as conventional murine strains are highly insensitive to these toxins. As a result, much of the pathogenesis work related to SAgs has been performed in rabbits (Salgado-Pabón et al., 2013; Tuffs et al., 2017; Wilson et al., 2011). Importantly, we have demonstrated that transgenic mice expressing the human leukocyte antigen (HLA)-DR4 are significantly more sensitive to SAg activity (Xu et al., 2015, 2014). This allowed us to determine that staphylococcal SAgs are important during bloodstream infections and also identified the liver as a key target for these factors (Xu et al., 2014).

In the current study, we deployed targeted antibody depletion protocols that demonstrated, during bloodstream infection, SAgs target CD4+ T cells to produce pathogenic levels of the key cytokine interferon-gamma (IFNγ). IFNγ promoted enhanced disease severity and bacterial burden in the liver and excess IFNγ levels during infection appeared to perturb liver macrophage activity to promote the survival of *S. aureus* within these cells. This the first report of targeted SAg activity that manipulates host macrophages to support *S. aureus* growth during bloodstream infections.

## RESULTS

### Transgenic HLA-DR4 C57BL/6 mice are sensitive to SAgs SEB and SEC and can model *S. aureus* bacteremia

Previously, we demonstrated that *S. aureus* burden is promoted during murine bloodstream infections by the SAgs staphylococcal enterotoxin A (SEA) and staphylococcal enterotoxin-like W (SElW); however, the mechanism remained uncharacterized (Vrieling et al., 2020; Xu et al., 2014). In the current study, we first utilized strain COL, a well-studied methicillin resistant *S. aureus* (MRSA) isolate from clonal complex (CC) 8 that produces the SAg, staphylococcal enterotoxin B (SEB). Splenocyte analysis from C57BL/6 (B6) or transgenic HLA-DR4 C57BL/6 (DR4-B6) animals, identified that T cell activation (measured by the production of IL-2) to titrating doses of SEB was orders of magnitude higher from spleen cells from the DR4-B6 animals compared with conventional B6 mice (Fig 1A). In addition, analysis of stimulated splenocytes using cytometry analysis demonstrated a massive expansion of Vβ8+ T cells, the major target of SEB in mice (Rellahan et al., 1990) (Fig 1B). These cells represent ∼20% of the T cell repertoire in the DR4-B6 animals and the majority of these were activated by SEB as measured by the upregulation of CD25 (Fig 1B). Together these data demonstrate that, compared to conventional B6 mice, DR4-B6 mice are highly sensitive to SEB and can be used for the analysis of SEB activity *in vivo*.

**Figure 1.**
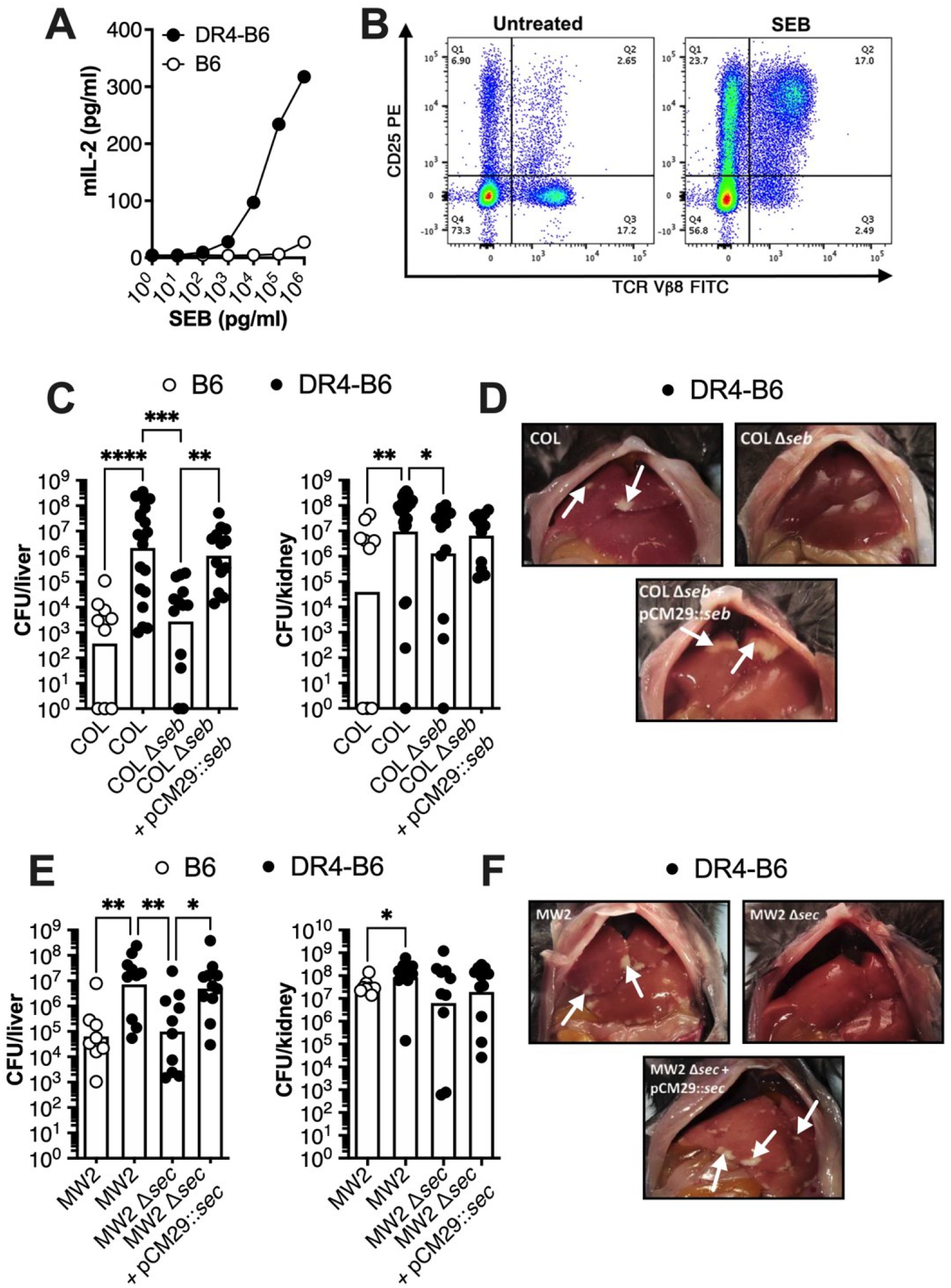
Superantigens SEB and SEC are important in *S. aureus* bacteremia when performed in transgenic HLA-DR4 C57BL/6 animals. (A) IL-2 production of isolated splenocytes from conventional C57BL/6 (open dots) and transgenic DR4-B6 (solid dots) mice following stimulation with a titration of SEB protein. (B) Activation of Vβ8+ T cells in DR4-B6 mice stimulated by SEB compared to the no-protein control as determined by CD25 expression (Quarter values represent total cell population). C57BL/6 and DR4-tg animals were inoculated intravenously (i.v.) with 5×10^6^ CFUs and then sacrificed at 96 hpi for *S. aureus* COL (C-D) and 72 hpi for *S. aureus* MW2 (E-F). Liver and kidney bacterial burden (C&E) was assessed in conventional B6 mice (open dots) or in transgenic DR4-B6 mice (solid dots). Each dot represents an individual mouse, and the bar indicates the geometric mean. Significant differences were determined using the Kruskal-Wallis test with uncorrected Dunn’s test for multiple comparisons (* p < 0.05, ** p < 0.01, *** p < 0.001, **** p < 0.0001). Representative livers from the infected mice from *S. aureus* COL and mutants (D) and *S. aureus* MW2 mutants (F), white arrows indicate the presence of liver lesions.

To determine if SEB contributes to pathogenicity in murine bacteremia, we infected B6 and DR4-B6 animals with *S. aureus* COL. We found that wild-type *S. aureus* COL was significantly more virulent in DR4-B6 mice with higher bacterial burden found in the liver and kidneys when compared to the B6 background (Fig 1C). This was due to SEB activity as the bacterial burden of the SEB-null mutant (COL Δ*seb*) essentially phenocopied the data obtained from the B6 animals. Importantly, this phenotype could be complemented with COL Δ*seb* containing pCM29::*seb* (Fig 1C and 1D). These data clearly demonstrate that SEB contributes to the pathogenicity of *S. aureus* COL during bloodstream infection.

To determine if additional SAgs other than SEB could also contribute to bacteremia, we expanded our analysis to include *S. aureus* MW2, a CC1 MRSA isolate that produces staphylococcal enterotoxin C (SEC) (King et al., 2016). SEC is phylogenetically similar to SEB, and has a similar Vβ activation profile in humans (King et al., 2016; Tuffs et al., 2018). We successfully deleted the SEC gene in *S. aureus* MW2 and were able to complement the gene *in trans* (Fig S1). Like *S. aureus* COL, we found a significant increase in bacterial burden in the liver and kidney in DR4-B6 animals compared to the B6 mice when infected with MW2 (Fig 1E). Furthermore, deletion of *sec* from MW2 resulted in a significant reduction in bacterial burden and liver pathology (Fig 1E & Fig 1F). These data indicate that both SEB and SEC, produced from different *S. aureus* backgrounds, can contribute to the pathogenesis of experimental bloodstream infection and that SAg-sensitive DR4-B6 mice are suitable for modelling SAg activity in the context of live *S. aureus* infection.

### Depletion of CD4+ T cells results in reduced bacterial burden in the liver of *S. aureus* infected DR4-B6 mice

It has been well-established that SAgs can target and activate different T cell subsets that express the appropriate TCR Vβ (Tuffs et al., 2018). For efficient nasopharyngeal infection, *Streptococcus pyogenes* required the expression of the SAg streptococcal pyrogenic exotoxin A (SpeA) (Kasper et al., 2014), and depletion of T cells resulted in a markedly reduced bacterial burden which phenocopied the deletion of the *speA* gene (Zeppa et al., 2017). In the current study, we used this T cell depletion strategy and applied it to our model of *S. aureus* bacteremia. We found that when CD4+ T cells were depleted, bacterial burden in the liver was significantly reduced, with a near complete reduction of visible lesions, while depletion of CD8+ T cells had no impact (Fig 2A). To reduce bacterial burden in the kidney, depletion of CD4+ T cells was not sufficient, and required the combined depletion CD4+ and CD8+ T cells to observe a phenotype, suggesting a limited role for CD8+ cells in this organ (Fig 2B). To determine if this effect was limited to conventional CD4+ T cells, we also depleted NK and iNKT cells using an anti-NK1.1 targeting antibody, according to a previously established protocol (Szabo et al., 2017a). In this case, there was no impact on bacterial burden or animal morbidity indicating these cells likely do not play a role in this infection model (Fig S2). Together these data indicate that bacterial burden in the liver during *S. aureus* bacteremia is promoted by the activity of conventional CD4+ T cells, likely due to activation by SEB.

**Figure 2.**
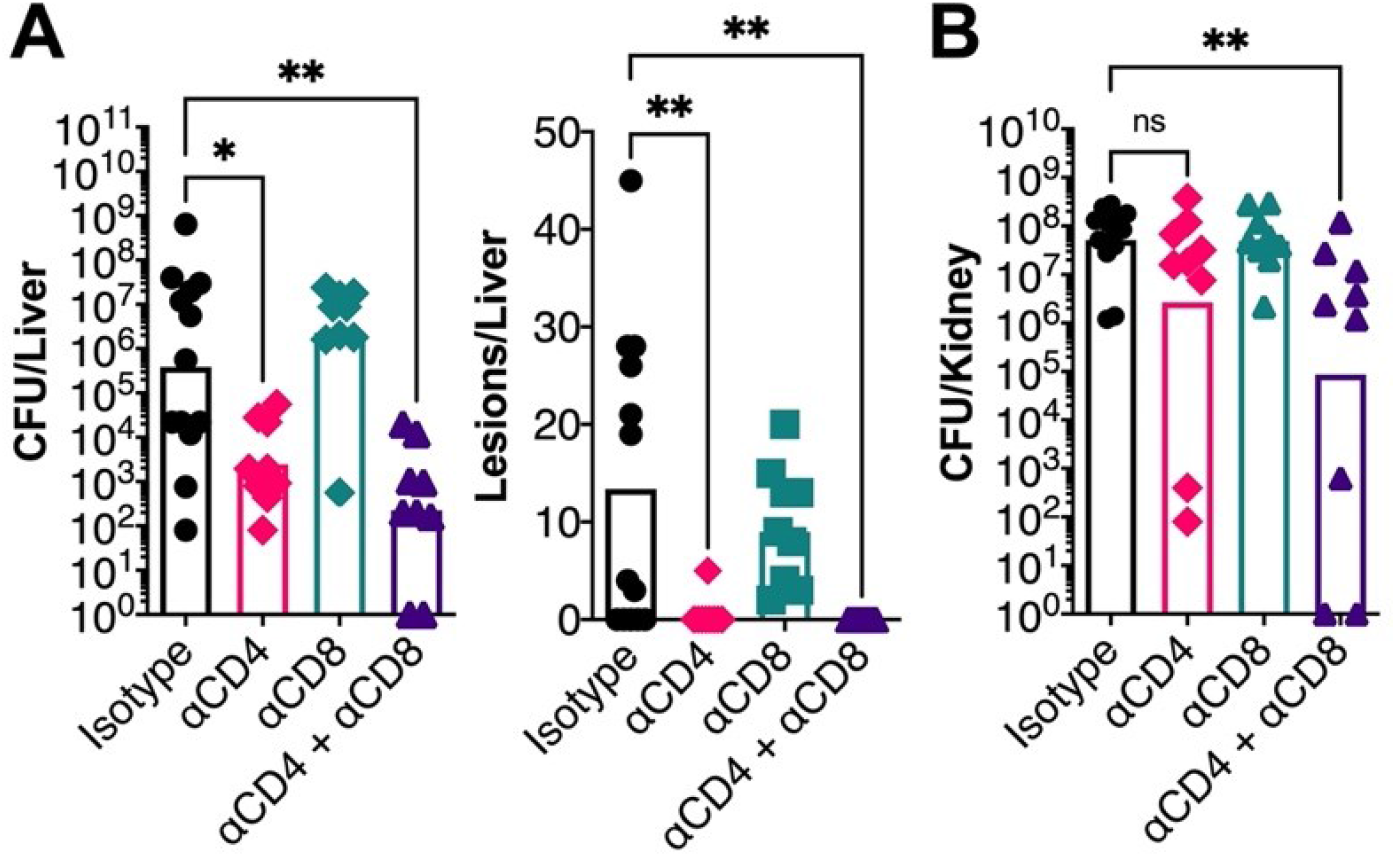
CD4+ T cells promote bacterial burden during *S. aureus* bloodstream infection. *In vivo* T cell depletion in DR4-B6 mice was performed with monoclonal antibodies to deplete CD4+ (clone GK1.5) and/or CD8+ (clone YTS169.4) cells prior to i.v. infection of *S. aureus* COL. *In vivo* liver bacterial burden and pathology (A), and kidney bacterial burden (B) was assessed 96 hpi. Each data point represents an individual mouse, and the bar indicates the geometric mean for CFUs/organ, and the median for lesions/organ. Significant differences were determined using the Kruskal-Wallis test with uncorrected Dunn’s test for multiple comparisons (* p < 0.05, ** p < 0.01).

### Blocking of IFNγ activity during systemic *S. aureus* infection results in reduced disease severity and bacterial burden

Previous analyses have demonstrated that CD4+ T cells can be targeted by staphylococcal SAgs to result in the release of numerous cytokines, including the key cytokines interferon-gamma (IFNγ) (also known as type II interferon), interleukin-10 (IL-10) and IL-17A (Tuffs et al., 2018). These cytokines have antagonistic activity to each other (Naundorf et al., 2009; Xu and Cao, 2010), and in the case of IL-17A and IFNγ, have been shown to contribute to SEB-mediated morbidity during toxemia models in HLA-transgenic mice (Szabo et al., 2017b; Tilahun et al., 2011). Together, this suggests that these key cytokines may contribute to SAg-mediated pathogenesis. To test this hypothesis, we used antibody depletion to block cytokine activity during bloodstream infection by both *S. aureus* COL and MW2 (Fig 3A). We found that only blocking of IFNγ resulted in a significant reduction in bacterial burden and liver pathology in the liver that phenocopied the deletion of *seb* or *sec* in *S. aureus* COL and MW2, respectively (Fig 3B and Fig 3E). Depletion of either IL-10 or IL-17A had limited impact on the liver burden suggesting that these cytokines do not promote bacterial burden in this model. Depletion of IFNγ also resulted in lower bacterial burden in the kidneys, suggesting that the blocking of this cytokine reduced the overall severity of this infection (Fig 3C and Fig 3F). It was also noted that bacterial burden in the kidney increased once IL-17A was depleted, but only for *S. aureus* COL. This suggests that IL-17A is important for protection against kidney damage during bloodstream infection by *S. aureus* in the HLA-transgenic mouse model. Overall, these data indicate that, of the three cytokines tested, only IFNγ promoted *S. aureus* burden during a bloodstream infection.

**Figure 3.**
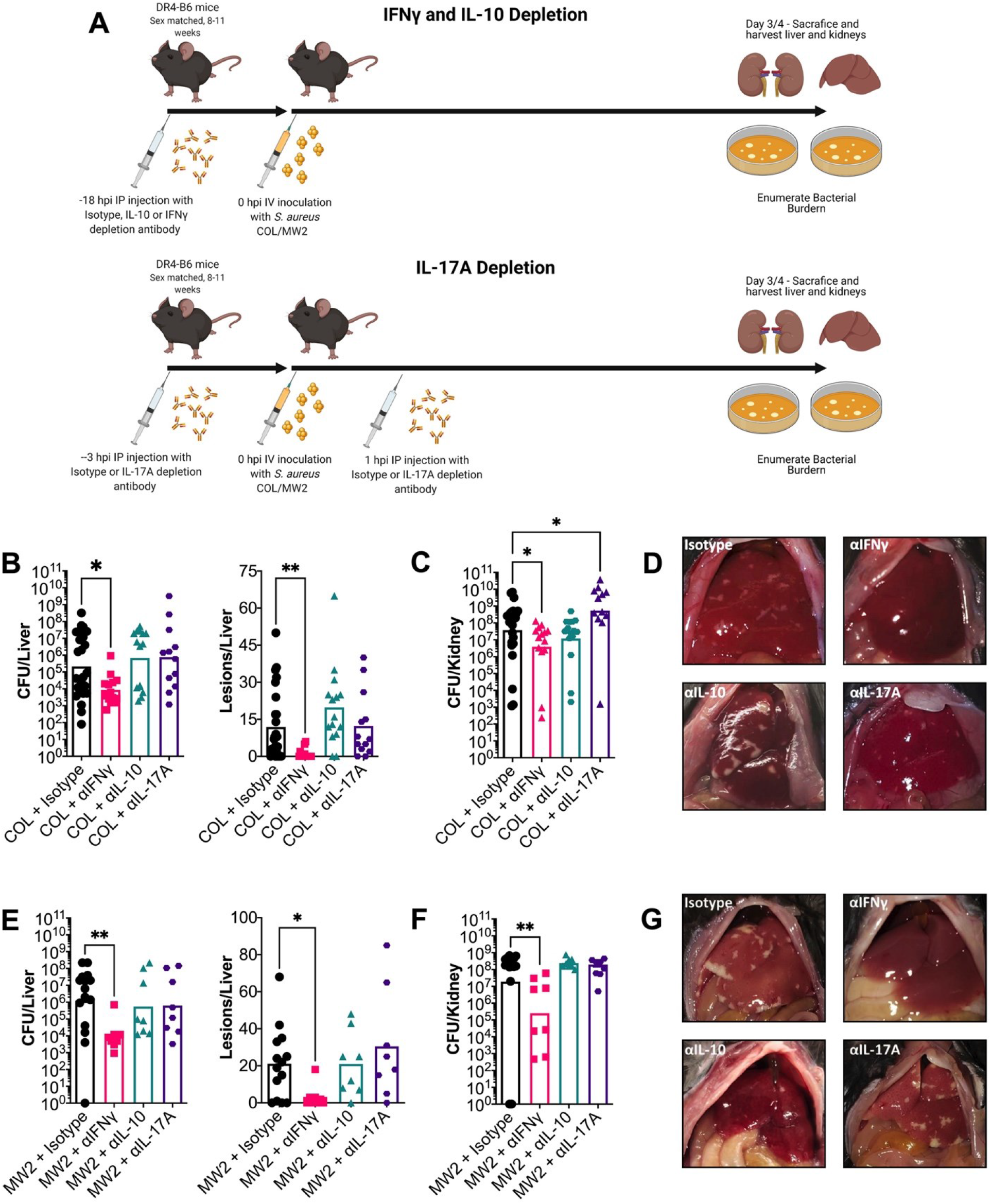
IFNγ promotes liver abscess formation and bacterial burden during *S. aureus* bacteremia in DR4-B6 mice. (A) Schematic outlining *in vivo* cytokine depletion with monoclonal antibodies prior to i.v. infection of DR4-B6 mice. Bacterial burden and abscess formation in liver (B&E) and kidney (C&F) at 96 hpi for *S. aureus* COL (B&C) or 72 hpi for *S. aureus* MW2 (E&F). Each dot represents an individual mouse, and the bar represents the geometric mean for CFUs/organ, and the median for lesions/organ. Significant differences were determined using the Kruskal-Wallis test with uncorrected Dunn’s test for multiple comparisons (* p < 0.05, ** p < 0.01). Representative livers from the cytokine treated mice from *S. aureus* COL (D) and *S. aureus* MW2 (F).

### Superantigens drive pathogenic production of IFNγ during *S. aureus* bloodstream infection

Several approaches were taken to determine if the promotion of bacterial burden by IFNγ was mediated by the SAg toxins. First, we performed cytokine analysis on liver homogenates and serum from animals infected with the wild-type or the SAg deletion mutants at 24 hours post infection (hpi). Compared to wild-type *S. aureus* COL infected mice, IFNγ levels were ∼10-fold lower in livers from animals infected with *S. aureus* COL Δ*seb* (Fig 4A). There was also a clear trend for more IFNγ in the serum for animals that were infected with the wild-type, although this did not reach statistical significance (Fig 4A).

**Figure 4.**
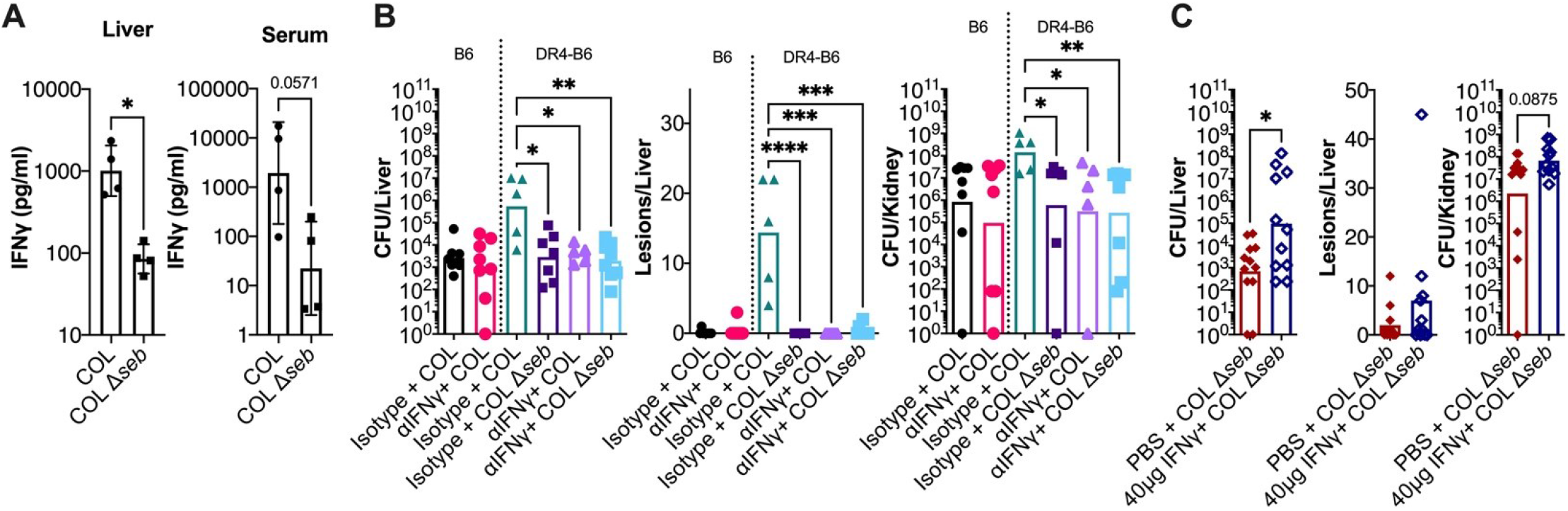
Superantigens promote pathogenic production of IFNγ that support bacterial burden. (A) DR4-B6 mice were inoculated i.v. with wild-type *S. aureus* COL or the COL Δ*seb* deletion strain. At 24 hpi animals were sacrificed and livers and blood were harvested, and material was prepared for IFNγ analysis. Each dot represents an individual mouse, the bar indicates the geometric mean, and error bars indicate the standard deviation. Dotted line indicated the levels detected in an uninfected animal. (B) C57BL/6 and DR4-B6 mice were treated 18 h prior to infection each animal was treated with 250 µg of isotype or αIFNγ antibody administered by i.p. injection. Animals were infected i.v. with *S. aureus* wild-type COL. (C) DR4-B6 animals were treated with 20µg (40µg total) of recombinant murine IFNγ or vehicle control (100 µl PBS) i.p. 2h before and 1h after i.v. infection with *S. aureus* COL Δ*seb*. In both experiments (B & C) i*n vivo* bacterial burden was assessed after 96 h in liver and kidneys and an assessment of gross pathological liver lesions was also performed. Each dot represents an individual mouse, and the bar indicates the geometric mean for CFUs/organ, and the median for lesions/organ. Significant differences were determined using the Mann-Whitney test (A, B & D) or Kruskal-Wallis test with uncorrected Dunn’s test for multiple comparisons (C) (* p < 0.05, ** p < 0.01, ***p < 0.001 ****p <0.0001).

Following from the cytokine analysis, we modified our infection model to characterise IFNγ depletion under circumstances where the SEB SAg from *S. aureus* COL was either absent or unable to function. In the SAg insensitive C57BL/6 background, bacterial recovery from infected mice was at a similarly low levels regardless of whether they had been treated with the IFNγ depletion antibody or isotype control, suggesting this phenotype can only be observed in a SAg sensitive environment (Fig 4B). Indeed, when we repeated the IFNγ depletion in the DR4-B6 background and included the COL Δ*seb* construct, bacteria recovered from organs of the isotype or IFNγ depleted groups were similarly low, whereas wild-type infections treated with the isotype control antibody produced visible liver lesions and higher bacterial counts in both the liver and kidney (Fig 4B). These data demonstrate that a functioning *seb* gene is required to promote pathogenic IFNγ activity.

Finally, to establish the link between SEB and IFNγ, we aimed to determine if the addition of exogeneous IFNγ could functionally complement the deletion of *seb* in *S. aureus* COL. In this experiment, animals were administered two 20 µg treatments of recombinant IFNγ 1 h prior to infection and 1 h after. Treatment with exogenous IFNγ resulted in a ∼2-log increase in bacterial burden in the liver when compared to the vehicle control (Fig 4C). Curiously, very few lesions formed on the surface of the liver with this approach, suggesting that sustained SAg/IFNγ activity is required for this pathology to become evident (Fig 4C). This demonstrates that the stimulation of pathogenic IFNγ is a key function of SEB during bloodstream infection and taken together with the previous data suggests a functional SAg must be present to elicit pathogenic production of IFNγ.

### IFNγ promotes early bacterial survival during *S. aureus* infection

With it established that SAgs could drive the production of pathogenic levels of IFNγ, we next wanted to determine when IFNγ had the most impact during the disease course. We therefore determined bacterial burden at shorter timepoints (i.e. 2 hpi, 8hpi, 12 hpi, 24 hpi, 36 hpi) in animals treated with αIFNγ antibodies or the isotype control (Fig 5A). From these data, much of the infectious dose became trapped within the liver following tail vein injection with ∼2×10^6^ CFU (approx. 40% of dose) at 2 hpi and this was followed by rapid clearance between 2 and 8 hpi in both groups. At 12 hpi the rate of bacterial clearance was reduced in the αIFNγ treated animals but continued steadily with almost complete clearance of the bacteria by 96 hpi. Conversely, after 24 hpi, in the livers of isotype treated, bacterial burden rapidly expanded reaching a level 3-logs higher by 96 hpi (Fig 5A). We performed a repeat of this analysis with daily timepoints and were able to confirm the trajectories that were observed in the shorter time-course (Fig S3). Together, these data indicate that IFNγ produced during wild-type infections by *S. aureus* is contributing to the ability of the bacteria to avoid clearance by the immune system in the liver during the early stages of bloodstream infection.

**Figure 5.**
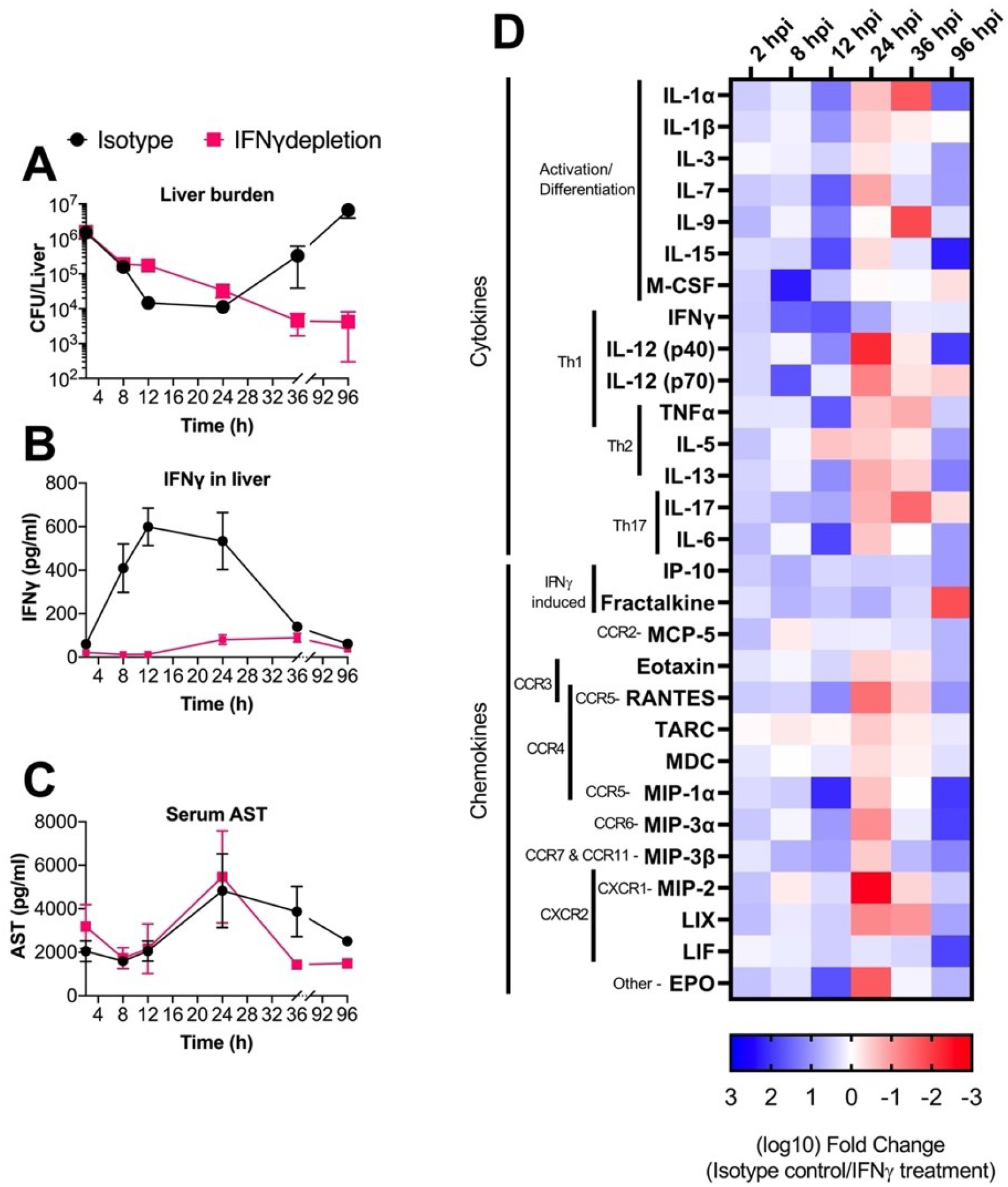
– SAg induced IFNγ promotes a pro-inflammatory environment that allows *S. aureus* to avoid complete clearance during bloodstream infection. Animals were treated with isotype or depleting IFNγ antibodies 18 h prior to infection with *S. aureus* COL. Following infection 3-4 animals were sacrificed from each group at the 6 timepoints shown, and liver and blood were harvested from each animal. Bacterial burden was determined at each timepoint and is shown as mean CFU/liver ± SEM (A). Liver homogenate was analysed by multiplex cytokine array and mean IFNγ (pg/ml) ± SEM at each timepoint were determined for each timepoint (B). Serum was analysed by ELISA to determine the concentration of Aspartate transaminase (AST), data shown are mean AST (pg/ml) ± SEM (C). Multiplex cytokine/chemokine array analysis was conducted on liver homogenate recovered from each animal sacrificed during the time-course (D). Data shown represent the log_10_ fold change for each cytokine between isotype and IFNγ depleted groups that displayed significant differences by students T test. Prior to comparison data was normalized to an antibody treated, uninfected animal sacrificed at the same timepoint as their comparator. Blue color (i.e. positive values) indicates more cytokine/chemokine is produced in the isotype infection and red color (i.e. negative values) indicates more cytokine/chemokine is produced in the IFNγ depleted infection. Broad classification for each cytokine is indicated, as well as potential binding partners for chemokines.

### Proinflammatory signalling is delayed and less intense when IFNγ is blocked during sepsis

IFNγ is a pleiotropic cytokine in the immune system and its depletion during infection could impact numerous downstream signalling pathways during the response to *S. aureus* bloodstream infection. In isotype treated mice, the highest level of IFNγ was observed in the liver between 12 and 24 hpi, in excess of 500 pg/ml (Fig 5B). Additionally, we confirmed that αIFNγ antibodies were able to reduce IFNγ concentration during infection up to 36 hpi (Fig 5B). We also analyzed serum samples for aspartate aminotransferase (AST) levels as a proxy for liver damage. These data indicate that there was limited change in liver damage during infection irrespective of IFNγ levels; however, there did appear to be a faster drop in AST levels in the IFNγ depleted groups in the later timepoints (Fig 5C), congruent with the reduced bacterial burden (Fig 5A).

To gain a broader understanding of the cytokine and chemokine dynamics, liver homogenates were analysed by multiplex cytokine array over the course of the experiment. This demonstrated that in earlier timepoints (2-12 hpi), many signalling molecules associated with inflammation were upregulated during infection where IFNγ was produced at high levels (Fig 5D). Strikingly, between 24 and 36 hpi, many cytokines and chemokines became reduced relative to the group treated with αIFNγ antibodies, which directly correlated with the expansion of *S. aureus* (Fig 5A) and was subsequently reversed again by 96 hpi (Fig 5D), suggesting that an inflammatory environment favourable to bacterial proliferation is sustained for a longer period during an infection where IFNγ production is high. Together, we infer that pathogenic production of IFNγ results in the rapid production of a pro-inflammatory environment in the liver that contributes to *S. aureus* survival during bloodstream infection.

### Macrophage activity in the liver is subverted by SAg-elicited IFNγ Production

Immune cells such as macrophages and neutrophils are critical for clearance of *S. aureus* during infection (Pidwill et al., 2021; Spaan et al., 2013). The cytokine and chemokine analysis indicated that wild-type *S. aureus* infection in HLA-DR4 mice drives an IFNγ-dependent pro-inflammatory signalling cascade in the livers of animals (Fig 5D). To determine if this response had any impact on the phagocytic cell populations in the liver, we first phenotyped immune cells isolated from this organ using flow cytometry at 24 and 96 hpi infection with *S. aureus* COL (see Fig S4 for gating strategy). We found few differences between the IFNγ-depleted or control mice in terms of the resident macrophages (Kupffer cells) (Fig 6A). For both monocytes and neutrophils, there were trends towards a higher percentage of these cells at 24 hpi in the isotype treated group, however, neither reached significance (Fig 6A). For neutrophils, this had subsided to a similar level in both groups by 96 hpi whereas monocytes had decreased in both groups by 96 hpi but there were significantly less in the IFNγ-depleted group at this time. We did detect significantly higher inflammatory macrophages in the IFNγ-depleted mice at 24 hpi although these were equivalent by 96 hpi (Fig 6A).

**Figure 6.**
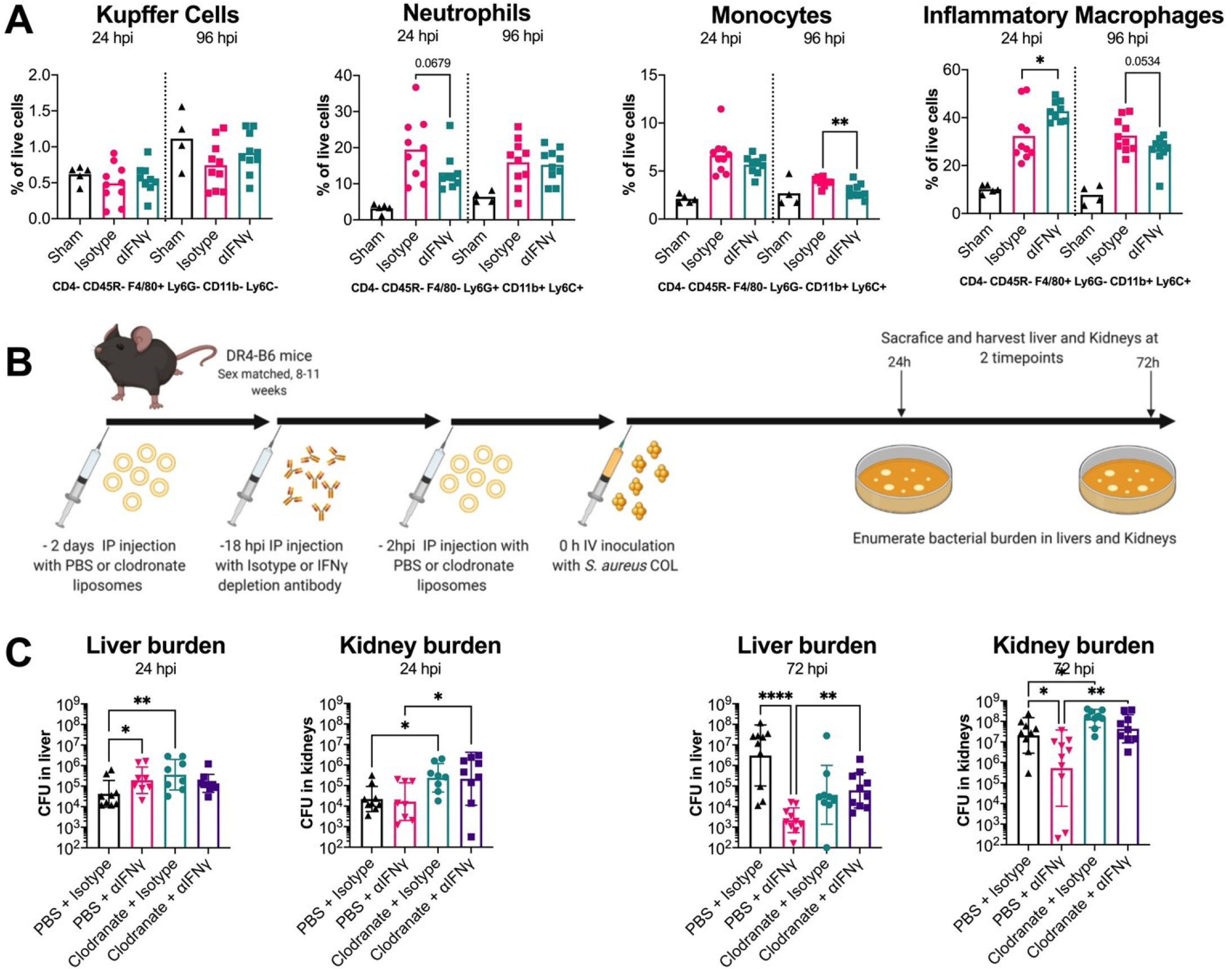
Liver macrophages are the target of pathogenic IFNγ production. (A) Flow cytometry-based phenotyping of immune cells isolate from livers infected of mice infected with *S. aureus* strain COL at 24 and 96 hpi. Control animals (sham) were treated with HBSS only. Cells were defined based on the staining profile listed below each graph and normalised to percentage of live cells. Istoype and αIFNγ treatments were compared, each dot represents an individual mouse, and the bar indicates the mean. Significant differences were determined using an unpaired Welch’s T test (* p < 0.05, ** p < 0.01). (B) Schematic outlining clodronate liposome-based depletion of macrophages along with IFNγ depletion used prior to i.v. infection of mice with *S. aureus* COL. (C) Bacterial burden in liver and kidney at 24 and 72 hpi are shown. Each dot represents an individual mouse, and the bar represents the geometric mean for CFUs/organ. Significant differences were determined using the Kruskal-Wallis test with uncorrected Dunn’s test for multiple comparisons (* p < 0.05, ** p < 0.01, ***p < 0.001 ****p <0.0001).

The infection kinetics (Fig 5A) and flow cytometry (Fig 6A) analysis suggest that the presence of high levels of IFNγ could impact the activity, recruitment, or differentiation of phagocytes, most likely inflammatory macrophages. To determine if these cells are the target of pathogenic IFNγ production, we depleted macrophages in mice using clodronate containing liposomes (Clodrosome®) and then performed *S. aureus* COL bloodstream infections with or without IFNγ blocking antibodies (Fig 6B). Macrophage depletion had an impact on animal welfare, so endpoints were brought forward from 96 hpi to 72 hpi and bacterial burden was determined in both the kidneys and the liver. In the liver, we again observed at 24 hpi, in animals treated with the control liposomes, that IFNγ depletion resulted in higher bacterial burden (Fig 6C). However, when we compared macrophage depleted animals the phenotype observed between the IFNγ depletion and isotype groups was abolished, indicating that macrophages are likely driving the IFNγ-dependent phenotype. We also observed an increase in bacterial burden when macrophages were depleted, compared to animals treated with control liposomes and isotype antibody (Fig 6C) indicating that these cells were important to restrict bacterial growth in the liver at this timepoint. In the kidneys, there was evidence that the depletion of macrophages likely resulted in greater bacterial ‘seeding’ to this organ although there was no difference due to IFNγ depletion (Fig 6C). The data from the 72 hpi timepoint confirmed the observation that macrophages were driving the IFNγ phenotype as again we were able to observe a clear IFNγ phenotype in both kidney and liver, yet this phenotype was mitigated by macrophage depletion (6D). Together, these data demonstrate that the promotion of *S. aureus* burden by IFNγ during bloodstream infection is mediated by macrophages.

### High levels of IFNγ allow for increased intracellular replication of *S. aureus* inside human macrophages

To determine if pathogenic IFNγ had any impact in the human system, white blood cells from healthy human donors were analysed for their responses to SAgs and IFNγ. First, we wanted to confirm that SAgs can elicit IFNγ from T cells through the engagement of the TCR. To do this, we compared IFNγ production elicited by recombinant SEB protein compared to the site directed mutant SEB-N23A. This mutant features a mutation within the TCR binding pocket resulting in a much lower ability to engage the TCR of its target cells (Leder et al., 1998). As expected, SEB-N23A elicited significantly lower IFNγ from human PBMC compared with wild-type SEB (Fig 7A). This confirmed that to elicit IFNγ from human PBMC, SEB must engage and activate the T cell through binding the TCR.

**Figure 7:**
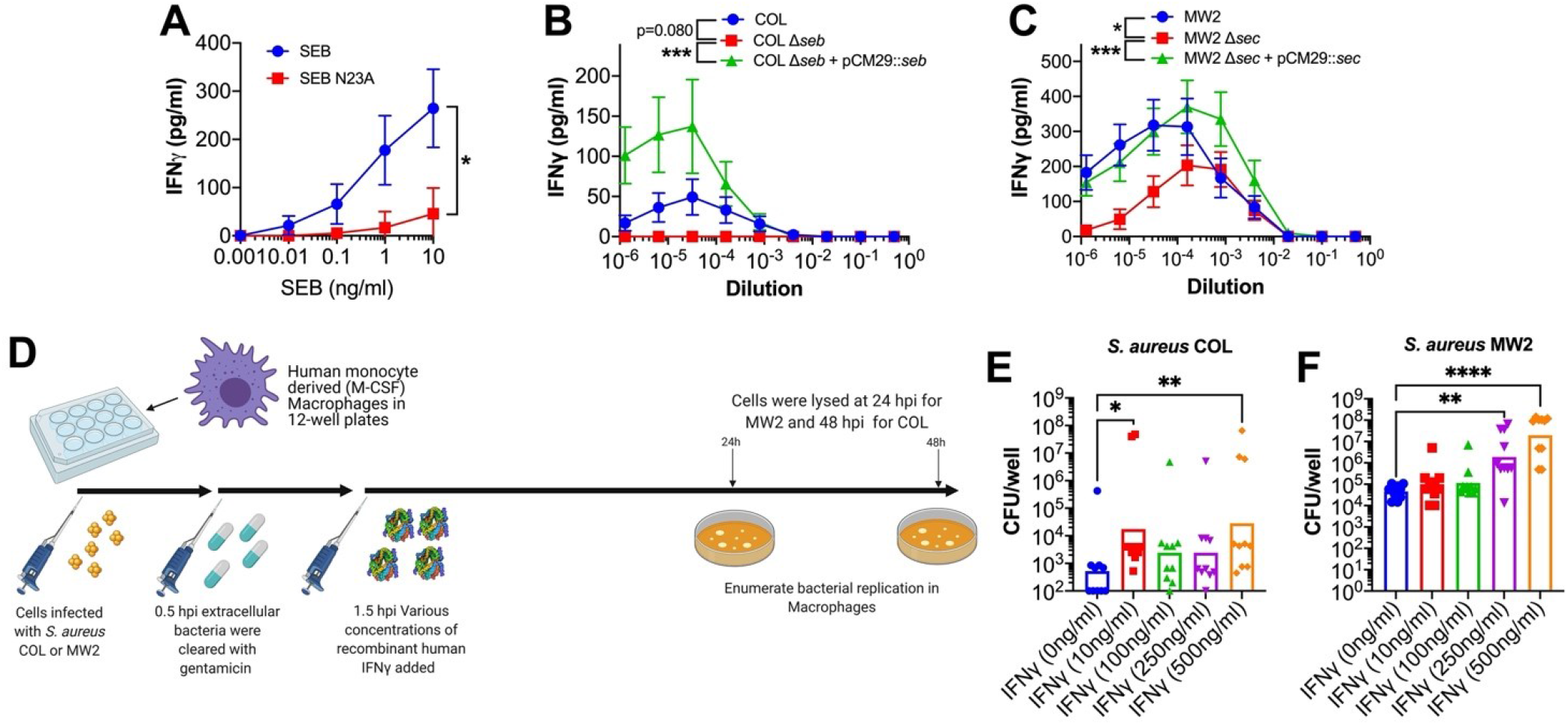
SEB and SEC elicit IFNγ from human cells and excessive concentration can promote increased extracellular replication of *S. aureus* in monocyte derived macrophages. (A) IFNγ production by PBMC from human blood following stimulation with a titration of SEB protein. IFNγ production by PBMC from human blood following stimulation with a titration of supernatant from *S. aureus* COL (B) or MW2 (C) constructs. Supernatants were taken from cultures grown for 8 h in BHI prior to use in these assays. Data shown (A-C) are mean ± SEM from 8 donors. Significant differences were determined from the area under each curve using a paired Friedman test for multiple comparison (* p < 0.05, *** p < 0.001). (D) Schematic outlining the procedure for intracellular *S. aureus* replication in monocyte derived human macrophages after dosing with recombinant human IFNγ. *S. aureus* recovered from human macrophages after incubation at 48 h for strain COL (D) and 24h for strain MW2 (E) with varying concentrations of recombinant IFNγ. Each dot represents macrophages form an individual human donor and the bar represents the geometric mean for CFUs/well. Significant differences between 0 ng/ml of IFNγ and other concentrations were determined using the Kruskal-Wallis test with uncorrected Dunn’s test for multiple comparisons (* p < 0.05, ** p < 0.01).

Next, we wanted to establish that our experimental strains (i.e. COL and MW2) could elicit IFNγ production from human T cells. We stimulated human PBMCs with a titration of wild-type bacterial supernatants grown for 8 h in brain heart infusion (BHI) broth and included supernatants from the respective SAg deletion and complemented strains. The data clearly indicated that both SEB and SEC, produced from COL and MW2 respectively, could drive IFNγ production in human PBMCs. The deletion of *seb* in COL eliminated the production of IFNγ, while there was a significant decline in the potency of MW2 Δ*sec*. The remaining IFNγ production was still easily detectable at lower MW2 Δ*sec* supernatant dilutions suggesting that other SAgs encoded by MW2 (i.e. *sea, selh, selk, sell, selq, selw* and *selx*) are also able to elicit the production of this cytokine.

As the murine model indicted macrophages are likely the major target of SAg-induced IFNγ, we infected human monocyte-derived macrophages with *S. aureus* and dosed these cells with varying concentrations of recombinant human IFNγ (Fig 7D). For both *S. aureus* COL and MW2, we saw an overall increase in intracellular bacterial replication when macrophages were dosed with high levels of IFNγ (Fig 7E and 7F). This phenotype was most evident with MW2 as this strain seems to have an improved ability at replicating inside macrophages. Moreover, replication was enhanced when IFNγ levels were high, with a nearly 2-log increase in bacterial burden when infected macrophages were treated with 500ng/ml of IFNγ (Fig 7F). Notably, the high concentrations of IFNγ did not impact macrophage viability and the bacteria were not simply overgrowing dead macrophages (Fig S5). Together, these data indicate that SAg-induced, IFNγ-mediated subversion of macrophages can occur in the human system, and that this mechanism appears to impair the ability of macrophages to kill intracellular *S. aureus*.

## DISCUSSION

Bloodstream infections caused by *S. aureus* represent a significant challenge in the clinic and SAgs have been shown to play a clear role in this disease (reviewed in Kwiecinski and Horswill, 2020; Spaulding et al., 2013). However, until now it has remained unclear how SAgs promote *S. aureus* persistence during infection (especially in the liver) and, specifically, how these toxins manipulate the response (Tuffs et al., 2018). The weak activity of SAgs in murine models has been a serious challenge in understanding how these toxins promote bacterial burden during infection. While rabbits are more sensitive to these toxins and may be more physiologically appropriate, challenges include both cost and the availability of advanced immunological tools (Salgado-Pabón and Schlievert, 2014). By using this HLA-transgenic mouse model, we can now examine the impact of the SAg-mediated cytokine response during *S. aureus* disease. By demonstrating that IFNγ can be promoted to pathogenic levels by the SAgs SEB and SEC, we provide the first report of a key cytokine produced in response to SAg-mediated T cell activation, that dramatically promotes bacterial burden during a systemic infection. This adds to a growing body of literature that demonstrates these toxins function to subvert different components of the immune system and challenges the waning dogma that these proteins are produced by the bacterium simply as an ‘immunological smoke screen’ (for a recent review discussing this see Tuffs et al. 2018).

The pathogenic potential of IFNγ has been alluded to in other studies. Firstly, a model of wound infection indicated that *S. aureus* capsular polysaccharide was shown to elicit IFNγ which led to an increased recruitment of neutrophils (McLoughlin et al., 2008). The authors postulated that higher neutrophil recruitment increased pathogenesis, as *S. aureus* was able to avoid neutrophil killing and survive intracellularly (McLoughlin et al., 2008). Our cytokine/chemokine data are consistent with these findings (Fig 5D), and while neutrophils were trending towards lower recruitment in the IFNγ depleted group at 24 hpi (Fig 6A), it appeared to be monocytes and inflammatory macrophages, rather than neutrophils in the isotype group that were favourably recruited to the liver by 96 hpi (Fig 6A). This variation in immune cell recruitment could promote a niche for the bacteria to reside within in the liver. Furthermore, our data also indicated IFNγ drove an increased production of the chemokine, fractalkine (CX3CL1), which has been suggested to be able to skew liver macrophages towards a more suppressive state (Aoyama et al., 2010). Added to this, high level of IFNγ itself was able to induce apoptosis and affect the life cycle of hepatocytes which can contribute to sustained inflammation in this organ (Horras et al., 2011). Together these observations, both in this study and others, suggest that SAgs through forcing the overproduction of IFNγ can modulate the liver environment to create a niche that is favourable for *S. aureus* survival.

The liver is an important organ during bacteremia as circulating pathogens are frequently filtered and trapped by resident macrophages (Kupffer cells) (Surewaard et al., 2016). In addition, there have been several reports that demonstrate *S. aureus* has evolved strategies to prevent this from occurring, including direct resistance to phagocytic killing by macrophages, or through the release of Hla that can aggregate platelets to create ischemic areas in the liver and promote further bacterial growth (Surewaard et al., 2016, 2018). It is important for the bacteria to establish in the liver as bacteria surviving here can eventually seed other organs, such as the kidney (Jorch et al., 2019; Surewaard et al., 2016).

Given the pleotropic nature of IFNγ, it is not surprising that several mechanisms may be at play in the liver to promote the growth of *S. aureus*. In addition to the other potential mechanisms of action defined in other studies, we have revealed a new pathway where macrophages are a clear target of pathogenic levels of IFNγ. To our knowledge, this is the first time that macrophage activity has been shown to be affected by high concentration of this cytokine to support intracellular replication of *S. aureus*. Of particular interest were the differences noted between the two *S. aureus* strains, while MW2 had the replicative ability to respond to IFNγ concentrations in a dose dependent manner, the same was not true for COL which featured considerable noise (Fig 7). Overall, these data suggest that COL as an isolate is not suited to replication within macrophages and may rely on another IFNγ driven mechanisms during these types of infections.

Our findings could also appear to be somewhat contradictory to several other studies that clearly demonstrate that IFNγ contributes to the clearance of *S. aureus* during bloodstream infection (Brown et al., 2015; Zhao et al., 1998). Furthermore, the activity of memory CD4+ T cells supports the clearance of *S. aureus* by producing this cytokine along with other signals to coordinate this response (Brown et al., 2015). These studies were conducted in conventional mouse strains that are less vulnerable to the activity of staphylococcal SAg and are more likely to represent what would occur in an immunocompetent individual, that is able to neutralise toxins like the SAgs. Indeed, this divergence is well presented by Brown et al. (2015), as in this study the IFNγ profiles of previously *S. aureus* exposed mice, demonstrate that this cytokine peaks almost immediately after infection and subsequently drops rapidly. For mice that were not pre-exposed to *S. aureus*, IFNγ was barely detected (Brown et al., 2015). This is contrary to what we have observed in SAg-mediated disease as IFNγ peaked later (24 hpi) and stayed high for much of the infection course. Together this suggests that IFNγ has a dual role during infection, primarily it is protective against *S. aureus* but if manipulated to high and sustained levels, can act as a mediator that promotes pathogenesis.

The implications of pathogenic IFNγ production in human health are significant. As discussed, *S. aureus* is one of the most common causes of bloodstream infection with disease often leading to life-threatening sepsis (Kwiecinski and Horswill, 2020). Indeed, sepsis is a very serious concern in the clinic, contributing to nearly 20% of global annual deaths (Rudd et al., 2020). One of the major challenges to treating sepsis is that without early intervention this disease can rapidly move from a microbiologically-mediated condition to an immunologically-driven sequela, often resulting in antibiotic treatment being ineffective (Corl et al., 2020). The pathophysiology of sepsis has also proven to be highly complex with many factors including the invading pathogen contributing to outcome. Death as an outcome of sepsis can occur both through acute inflammatory processes that leads to multi-organ failure as well as chronic immunosuppressive activity (Van Der Slikke et al., 2020). In both cases, IFNγ can contribute to these outcomes as a key promoter of the proinflammatory response, or due to its absence leading to the dominance of immunosuppressive pathways (Hotchkiss et al., 2013; Romero et al., 2010). Indeed, several studies have demonstrated that once a patient enters the immunosuppressive state of sepsis, therapy with IFNγ may actually improve outcomes, however, the opposite maybe true if administered too early (Nalos et al., 2012; Payen et al., 2019).

There is also evidence to suggest this mechanism may be at play in the context of *S. aureus* vaccines and could be an important consideration for vaccine design. Karauzum *et al* (2017) found that whole cell vaccines in mice promoted disease and bacterial survival, through the activity of a heavily skewed Th1 immune response. It appeared in this bloodstream infection model that disease was promoted by the vaccines and this was mediated by excessive production of IFNγ (Karauzum et al., 2017). It was also suggested that this study had significant parallels with the failure of a clinical trial using the IsdB subunit vaccine, which was intended to protect against bacteremia, but instead had to be terminated early as it worsened patient outcome (Fowler et al., 2013; Karauzum et al., 2017). Together this would suggest that *S. aureus* has evolved to take advantage of a human immune system whose responses have been skewed by the activity of IFNγ and further to this, evolved a family of toxins capable of driving this skewing itself.

In conclusion, we report the discovery that *S. aureus* SAgs, SEB and SEC, can drive the production of IFNγ during bloodstream infection to promote disease. Our analyses suggest that that the pathogenic production of IFNγ subverts macrophage activity allowing the bacterium to persist within the liver leading to increase morbidity. Furthermore, we were able to establish this mechanism has implications for human health as IFNγ can promote bacterial intracellular replication in human macrophages. Together this moves forward our understanding of the immunological factors at play during *S. aureus*-meditated sepsis in the context of pathogen-driven inflammation and can inform on appropriate design of treatments and vaccines targeting *S. aureus* disease.

## ACKNOWLEDGEMENTS

This work was supported by an operating grant from the Canadian Institutes of Health Research (CIHR) (PJT-166050) to S.W.T and J.K.M. D.E.H acknowledges funding from CIHR grant PJT-153308. Figure 3A, 6B and 7D were created in part using Biorender.com.

## AUTHOR CONTRIBUTIONS SECTION

S.W.T executed the majority of the experimental work assisted by M.I.G, S.X.X, H.C.C, K.J.K, J. C. and D.E.H. Experimental design and data interpretation were performed by S.W.T, M.I.G, S.M.K, R.S.F, D.E.H, and J.K.M. S.W.T and J.K.M conceptualized the study and wrote the manuscript, which was reviewed and approved by all co-authors.

## MATERIALS AND METHODS

### Human Ethics Statement

Human venous blood was taken from healthy donors in accordance with a human subject protocol approved by the London health sciences centre (LHSC) research ethics board, Western University, London, Ontario, Canada, under the protocol 110859. Volunteers were recruited by a passive advertising campaign within the Department of Microbiology and Immunology at Western University and following an outline of the risks, written informed consent was given by each volunteer before each sample was taken. Following sampling, blood was fully anonymized and no information regarding the identity of the donor, including sex and age, was retained.

### Mice

Eight-to-eleven-week-old male and female HLA-DR4-IE (DRB1*0401) humanized transgenic mice lacking endogenous mouse MHC-II on a C57BL/6 (B6) background (here referred to as DR4-B6 mice) (38) or B6 mice were used for all *in vivo* infection experiments. DR4-B6 animals were bred onsite at Western University and B6 mice were purchased directly from the Jackson Laboratory (Stock N° 000664). Animals for experiments were housed in single sex cages which did not exceed 4 in number. During all breeding and experiments, mice were provided food and water *ad libitum* and appropriate enrichment was provided in all cages. All animal experiments were in accordance with the Canadian Council on Animal Care Guide to the Care and Use of Experimental Animals, and the animal protocol was approved by the Animal Use Subcommittee at Western University.

### Bacterial strains, media, and growth conditions

*S. aureus* strains listed in Table S1 were grown aerobically at 37°C in tryptic soy broth (TSB) (Difco) or brain heart infusion broth with shaking (250 rpm) supplemented with the appropriate antibiotics. For solid phase cultures tryptic soy agar (TSA) was used (TSB+ 1.5% w/v agar, Fisher Scientific)) supplemented with the appropriate antibiotics. *Escherichia coli* strains were used as cloning hosts and were grown in Luria-Bertani (LB) broth (Difco) or LB agar supplemented with appropriate antibiotics at 37°C with shaking (250 rpm). Growth curve analysis was performed using a Biotek Synergy H4 multimode plate reader.

### Construction of MW2 Δ*sec* mutant

Markerless deletion of *sec* in MW2 was performed using the pKor1 allelic replacement system (Bae and Schneewind, 2006). Briefly, a 598 bp fragment upstream of *sec* was amplified with Phusion™ High-Fidelity DNA Polymerase (Thermo Fisher) using the primers (Table S2) pKOR-sec-upstream-For and pKOR-sec-upstream-Rev along with a 576 bp region downstream of *sec* amplified by the primers pKOR-sec-downstream-For and pKOR-sec-downstream-Rev 2. These products contained a 12 bp overlap and were spliced together at a ratio of 1:1 using primers pKOR-sec-upstream-For and pKOR-sec-downstream-Rev, creating an insert of 1203 bp in total. This insert was integrated into empty pKOR1 using BP clonase (Thermo Fisher) according to the manufacture’s instructions. The cloned plasmids were transformed into *E. coli* XL1-Blue and screened for plasmids containing the insert. The confirmed knockout construct was chemically transformed into *E. coli* SA30B (Monk et al., 2015) to methylate the plasmid for electrotransformation into *S. aureus* MW2 (Monk et al., 2012). The *sec* knockout was created as described previously (Bae and Schneewind, 2006) and candidate constructs were screened by PCR using primers SEC-screen-For and SEC-screen-Rev (Table S2).

### Construction of pCM29::*seb* and pCM29::*sec* complementation plasmids

SEB and SEC-complementation plasmids for *S. aureus* SAg null mutants were created as previously described, with modifications (Vrieling et al., 2020). Briefly, SAg coding sequences were cloned into a pCM29 vector containing the active promotor of the leukocidin LukMF’ (Vrieling et al., 2015). To achieve this, pCM29::pLukM-sGFP was digested with KpnI and EcoRI to remove the sGFP coding sequence while retaining the *lukM* promotor sequence. SAg insert fragment forward primers were designed to contain endogenous RBS upstream of the start codon as this would be removed from the plasmid with the *sgfp* gene. Sequences of *seb* and *sec* were amplified respectively from COL and MW2 genomic DNA using Phusion™ High-Fidelity DNA Polymerase (Thermo Fisher) with primers listed in Table S2. PCR products were digested with KpnI and EcoRI to prepare the SAg insert for ligation. Complementation inserts were ligated into pCM29 that had the *sgfp* removed with T4 ligase (NEB). Following ligation, plasmids were further digested with MluI to inactivate any contaminating pCM29 that still retained the *sgfp*. After this step, ligations were transformed into *E. coli* SA30B (Monk et al., 2015) for appropriate methylation before transformation, of sequence positive constructs, into electrocompetent *S. aureus* using a protocol previously described (Monk et al., 2012).

### Protein expression analysis

Recombinant staphylococcal enterotoxin B (SEB) was generated as described previously (Chau et al., 2009). Briefly, SEB was expressed with a His-tag in BL21 (DE3) *E. coli* and purified by nickel column chromatography. An attenuated mutant of SEB that has impaired binding to TCR was also purified. The mutant SEB carries an N→A point mutation at position 23 and is referred to as SEB_N23A_ (Hayworth et al., 2012; Leder et al., 1998).

### Murine splenocyte analysis

The ability of murine cells to respond to SEB was determined using interleukin-2 (IL-2) production. Mouse spleens were removed and broken into a single-cell suspension, followed by red blood cell lysis in ammonium-chloride-potassium (ACK) buffer. The remaining cells were suspended in complete RPMI (cRPMI), containing RPMI (Invitrogen Life Technologies) supplemented with 10% Fetal Bovine Serum (FBS) (Wisent Inc., Quebec, Canada), 100 μg/ml streptomycin, 100 U/ml penicillin (Gibco), 2 mM l-glutamine (Gibco), 1 mM sodium pyruvate (Gibco), 100 μM nonessential amino acids (Gibco), 25 mM HEPES (pH 7.2) (Gibco), and 2 μg/ml polymyxin B (Gibco). Cell suspension was seeded into 96-well plates at a density of 1.1 × 10^6^ cells/ml. Titrating concentrations of recombinant SEB were added to cells and incubated for 18 h at 37°C with 5% CO2. Supernatants were assayed for IL-2 by enzyme-linked immunosorbent assay (ELISA) according to the manufacturer’s instructions (Thermo fisher). For flow cytometry, cells were dual stained with phycoerythrin (PE)-conjugated anti-CD25 (clone PC61.5) (eBioscience) and FITC-conjugated anti-Vβ8 (clone KJ16) (eBioscience). Events were acquired using a FACSCanto II (BD Biosciences), and data were analyzed using FlowJo v.10.7.1 TreeStar).

### Staphylococcal bacteraemia model

Single bacterial colonies were picked from a TSA plate and grown in 3 ml TSB overnight (16 to 18 h). Cells were subsequently subcultured in TSB to an OD_600_ of 0.1 and grown to post-exponential phase (OD_600_ ∼3.0 to 3.5). The bacterial pellet was washed once and resuspended in HBSS to an OD_600_ of 0.15 for strain COL and 0.85 for strain MW2, corresponding to ∼ 5 × 10^7^ CFU/ml. Mice were injected via the tail vein with 5 × 10^6^ CFU of *S. aureus* in a total volume of 100 μl. Mice were weighed and monitored daily. At various timepoints post-infection, mice were sacrificed (maximum of 3 and 4 days for MW2 and COL, respectively), and the kidneys and liver were aseptically harvested. All organs were homogenized, plated on mannitol salt agar (Difco), and incubated at 37°C overnight. *S. aureus* colonies were enumerated the following day with a limit of detection determined to be 3 CFU per 10 μl.

### Antibody depletion protocols

CD4+ and CD8+ T cells were depleted in animals according to a protocol described previously (Zeppa et al., 2017). Briefly, mice were injected with 300 µg of T-cell depleting antibodies [anti-CD4 (clone GK1.5, BioXCell); anti-CD8 (clone YTS169.4, BioXCell); or both, at 150 µg each] or isotype control (clone LTF-2, BioXCell) 7, 6 and 1 day before infection with *S. aureus*. For IL-17A depletion, mice were treated with 200 µg dose of an anti–IL-17A mAb (clone 17F3; BioXCell,) or a mouse IgG1 isotype control (clone MOPC-21, BioXCell) 3 h before *S. aureus* infection, then with a further 100 µg dose 1 h after infection, as described previously (Szabo et al., 2017c). For IL-10 and IFNγ depletions, mice were treated with a 250 µg dose of anti-IL-10 mAb (clone JES5-2A5), anti-IFNγ (clone XMG1.2, BioXCell) or Rat IgG1 isotype control, anti-horseradish peroxidase (clone HRPN) 18 h prior to infection. NK cells were depleted in mice with 200 μg of anti-NK1.1 mAb (clone PK136, BioXCell) or mouse IgG2a isotype control (clone Cl.18, BioXCell) administered 18 h prior to infection, as described previously (Hayworth et al., 2012). All antibody doses were prepared in 100µl – 200 µl PBS and administered by intraperitoneal (i.p.) injection.

### Detection of cytokines and chemokines *in vivo*

At various time point post-infection, serum supernatants and livers were collected. Supernatants were obtained from whole livers by homogenization in HBSS supplemented with the complete protease inhibitor cocktail (Roche). Samples were analyzed using Mouse Cytokine Array/Chemokine Array 44-Plex (MD44, Eve Technologies). AST levels were assessed from murine serum using a mouse aspartate aminotransferase (AST) ELISA Kit (Abcam).

### Flow cytometry analysis of murine cells

Livers were extracted from mice and pushed through a 0.7 µm cell strainer. Leukocytes were isolated from livers using a 33.75% Percoll gradient (GE Healthcare). Following isolation, red blood cells were lysed using ammonium-chloride-potassium (ACK) lysis buffer (Gibco) and washed with PBS containing 2% FBS. Cell viability was first determined using Fixable Viability Dye eFluor™ 506 (Thermo Fisher) and then subsequently stained anti-CD4-PE-Cy5 (clone RM4-5, Thermo Fisher) anti-CD45r-V450 (clone RA3-6B2, BD), anti-F4/80-A647 (clone BM8, Biolegend), anti-Ly6G-A700 (clone RB6-8C5, Biolegend), anti-Ly6C-BV711 (clone RB6-8C5, Biolegend), and anti-CD11b-PE (clone M1/70, Biolegend). Cells were fixed overnight with 1% paraformaldehyde prior to analysis. Events were acquired and data analyzed as outlined above. Events were acquired using a LSR II (BD Biosciences), and data were analyzed using FlowJo v10.7.1 (TreeStar).

### Macrophage depletion in mice

Macrophage depletion was based on a protocol previously described (Stritzker et al., 2010). Briefly, 200 µl of Clodronate containing liposomes and control liposomes [Clodrosome® + Encapsome® (Encapsula Nano Sciences)] were administered to the mice i.p. 2 days and 4 h prior to infection with bacteria. At 18 h prior to infection, IFNγ depleting or control antibodies were also administered to the mice.

### Detection of human cytokines from stimulated human cells

The ability of human cells to produce cytokines was determined from stimulated peripheral blood mononuclear cells (PBMC). These cells were isolated from human blood by density-based centrifugation following layering of the blood onto Ficoll-Hypaque plus (GE healthcare). Cells were isolated and washed three times in RPMI (Gibco) and then resuspended in cRPMI. Cell suspension was seeded into 96-well plates to a final concentration of 1.0 × 10^6^ cells/ml. Titrating concentrations of recombinant proteins or *S. aureus* supernatants were added to cells and incubated for 18 h at 37°C with 5% CO_2_. Supernatants were assayed for IL-2 or IFNγ by enzyme-linked immunosorbent assay (ELISA) according to the manufacturer’s instructions (Thermo fisher).

### Human macrophage cultures and infections

Primary human macrophages were derived from blood monocytes isolated from healthy human volunteers as previously described (Flannagan et al., 2012, 2016). Briefly, mononuclear cells were isolated from blood with lympholyte®-poly (Cedarlane Laboratories) according to the manufacturer’s instructions. Monocytes adhered to glass coverslips in 12-well plates (1.5×10^6^ cells/well) and were subsequently cultured for 7–9 days in RPMI (Gibco) with 10% FBS (Wisent) and 0.5 ng/ml recombinant human Macrophage Colony Stimulating factor (M-CSF) (R&D Systems) to allow for differentiation of monocytes into macrophages. After 5 days of differentiation, adhered cells were washed with PBS, and the medium was replaced with fresh RPMI + 10% FBS containing M-CSF. Macrophages were differentiated to day 7 and used experimentally until day 10.

*S. aureus* strains COL and MW2 were cultured overnight in TSB then pelleted and re-suspended in serum free RPMI and then diluted in serum free RPMI to an OD_600_ of 0.5. Cells were infected with an MOI of 30 and following inoculation were centrifuged at 277 × g for 2 min, then incubated for 30 min at 37°C in the presence of 5% CO_2_. Following phagocytosis, cells were treated with RPMI containing gentamicin (100 µg/mL) for 1 h at 37°C to kill extracellular bacteria. After gentamicin treatment, macrophages were rinsed with PBS and incubated further in RPMI containing 10% FBS without antibiotic. At this point, recombinant human IFNγ (R&D Systems) was also added at varying concentrations. Macrophages were incubated for 24 or 48 h following infection with MW2 or COL, respectively. Enumeration of antibiotic-protected bacteria (i.e. intracellular bacteria) was performed by lysing infected macrophages with 0.1% (v/v) Triton X-100 in PBS. Macrophage lysates were serially diluted and plated on TSA for enumeration.

### Statistical analysis

All statistical analysis were performed using GraphPad prism 9. In all tests a P value < 0.05 was considered statistically significant. For all bacterial burden CFU, analysis was performed with non-parametric Mann Whitney or Kruskal-Wallis test with uncorrected Dunn’s test for multiple comparisons, depending on group numbers. Flow Cytometry data was analysed using Welch’s T test to determine significant differences between means of the isotype and IFNγ depleted groups.

For the multiplex cytokine analysis heat map shown in Fig 5D, each raw data point was normalized to a sample taken from an uninfected control animal that was sacrificed at the same timepoint after treatment with the same antibody. Following normalization, the data from analysis at each timepoint was compared for statistical significance between the Isotype and IFNγ depletion using a students T test. All significant values were extracted, and the mean quantity of the cytokine/chemokine detected in the isotype treated animals was divided by the quantity detected in the IFNγ depleted group. These values were converted to log_10_ values to give fold-change in positive and negative values that could be plotted on a heatmap.

For human cytokine analysis performed in Fig 7A-C, the area under each curve for each donor was determined. These values where then compared using a paired T test or paired Friedman test for multiple comparisons, depending on the number of groups. Paired tests were used due to the large variation observed between individual human donors.

