## Supplemental figures and tables for "Superantigens promote *Staphylococcus aureus* bloodstream infection by eliciting pathogenic interferon-gamma (IFNγ) production that subverts macrophage function"

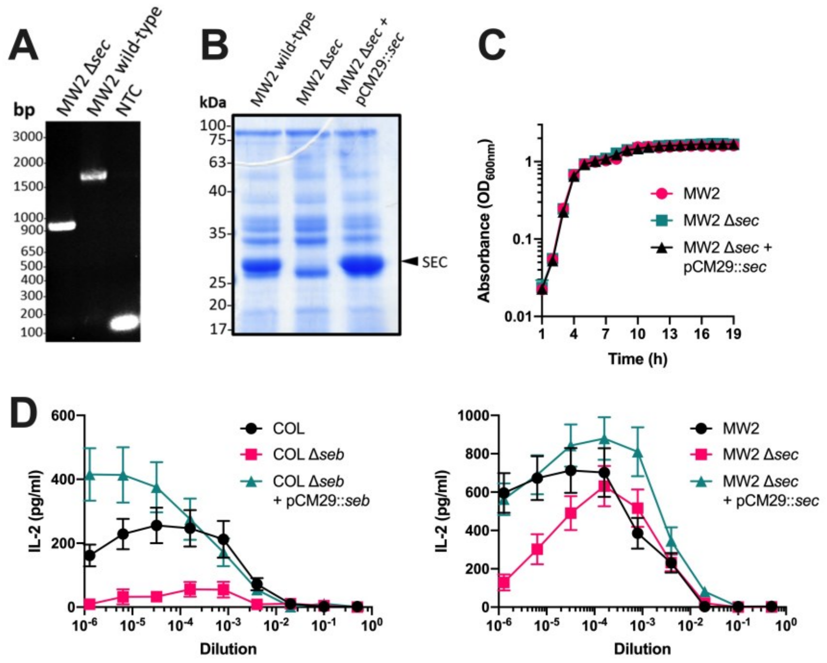


**Figure S1. Validation of *S. aureus* COL and MW2 superantigen mutants.** (A) Following allelic replacement, the deletion of *sec* gene was confirmed by PCR using primer SEC-screen-For and SEC-screen-Rev (Table S1). PCR products were run on a 1% gel and a PCR with genomic DNA from wild-type MW2 was included for comparison, along with a no template control (NTC). (B) *S. aureus* MW2 strains were grown in BHI broth for 8 h and the secreted profile was assessed by SDS PAGE analysis on 12% acrylamide gel stained with instant blue (Coomassie based stain). (C) To ensure that the *sec* deletion in MW2 had no impact on general viability, growth curve analysis was performed on the MW2 strains. Each clone was grown in TSB over the course of a 19 h with measurements taken every hour. Data shown are mean ± SEM. (D) To confirm the ability of these constructs to be able to activate T cells, IL-2 production was determined. PBMC from human blood where isolated and stimulated with a titration of supernatant from *S. aureus* COL or MW2 strains. Supernatants were taken from cultures grown for 8 h in BHI prior to use in these assays. Data shown are mean ± SEM from 8 donors.

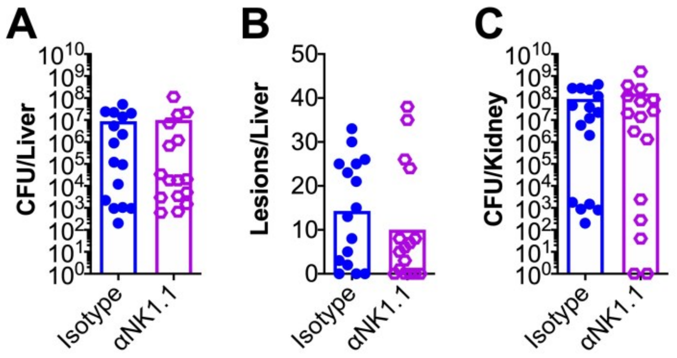


**Figure S2. Depletion of NK1.1+ cells does not alter *S. aureus* burden or pathology during bloodstream infections.** *In vivo* depletions in DR4-B6 mice were performed with monoclonal antibodies to deplete NK1.1 cells prior to intravenous infection of *S. aureus* COL. *In vivo* liver bacterial burden (A), liver pathology (B), and kidney bacterial burden (C) was assessed 96 h post i.v. challenge. Each data point represents an individual mouse, and the bar indicates the geometric mean for CFUs/organ, and the median for lesions/organ. No significant differences were detected.

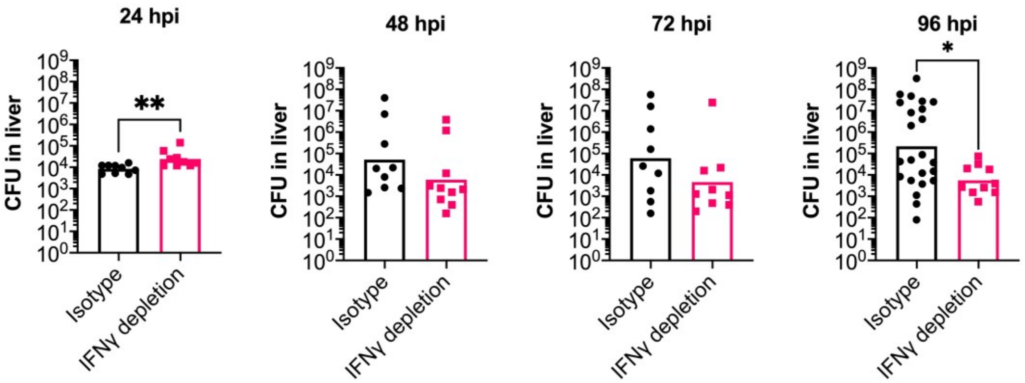


**Figure S3. Daily Time-course of *S. aureus* liver burden during bloodstream infection.** Animals were treated with the isotype control or IFNγ-depleting antibody 18 h prior to infection with *S. aureus* COL. Following infection, animals were sacrificed from each group at the 4 timepoints show, and livers were harvested from each animal and bacterial burdens determined. Each dot represents an individual mouse, and the bar indicates the geometric mean. Significant differences were determined using the Mann-Whitney test (* $p < 0.05$, ** $p < 0.01$,). For comparison, data shown in the 96 hpi timepoint are the same data included in Fig 3b (lacking the IL-17A depletion and isotype controls).

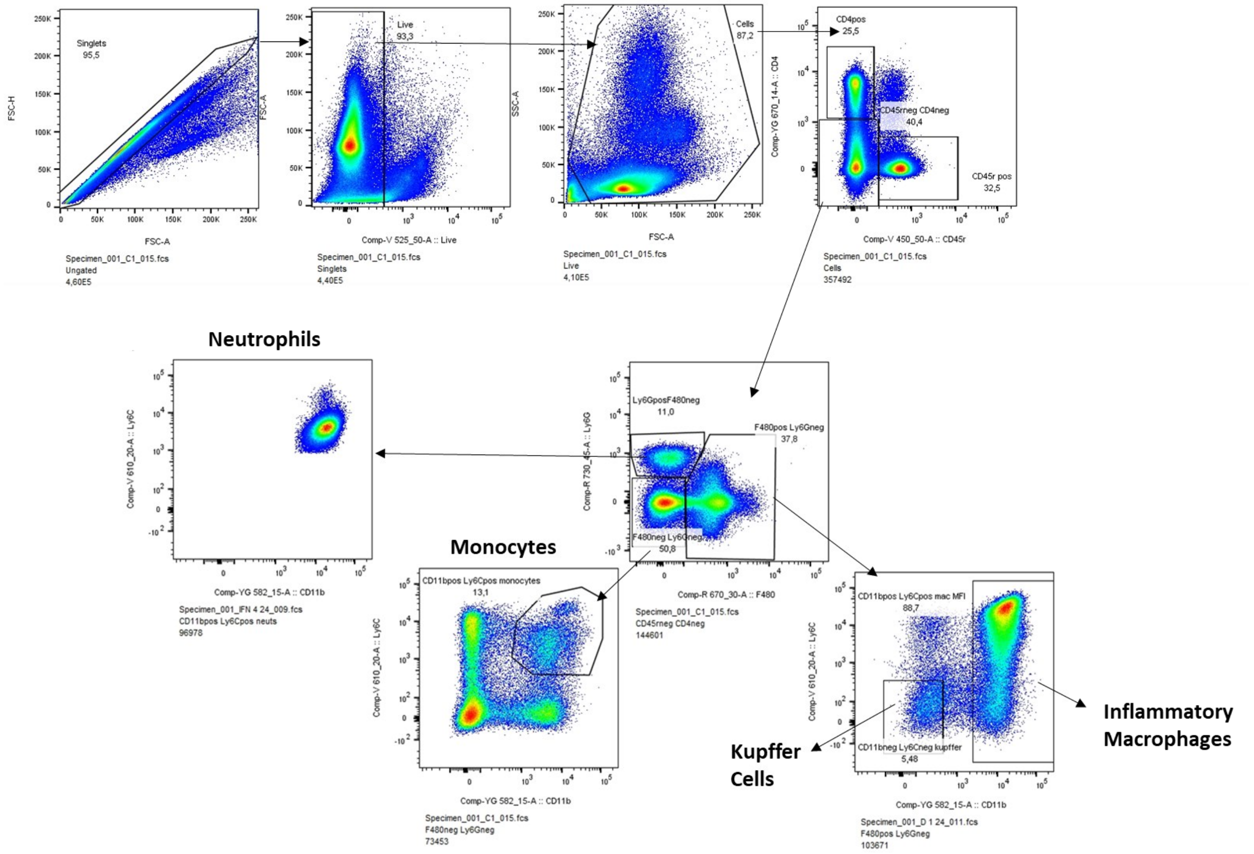


**Figure S4. Flow cytometry gating strategy for liver analysis.** Following isolation from murine livers, immune cells were analysed by flow cytometry. Prior to cytometer analysis cells were stained with antibodies against F4/80, CD11b, Ly6C, Ly6B, CD45r and CD4. Also included was a live/dead stain to determine cell viability. Schematic of cell phenotyping is shown with arrows denoting the workflow for gating. Cells were gated first on whether they were liver or dead and the live population was subsequently gated for the singlet population. Single liver cells were checked for the expression of CD4 and CD45r markers and the double negative population (i.e. majority myeloid derived cells) were gated for further analysis. Cells were then checked for their expression of F4/80 and Ly6G. Cells that were subsequently high for Ly6G but negative for F4/80 were checked for CD11b and Ly6C expression and classed as neutrophils. Ly6C and F4/80 double negative cells were gated and checked for CD11b and Ly6C expression, with double positive cells classed as monocytes. F4/80 positive and ly6G negative cells were checked for CD11b and Ly6C expression. Cells that were negative for both were considered resident macrophages i.e., Kupffer cells. Cells remaining cells in this group that expressed high levels of CD11b were considered inflammatory macrophages.

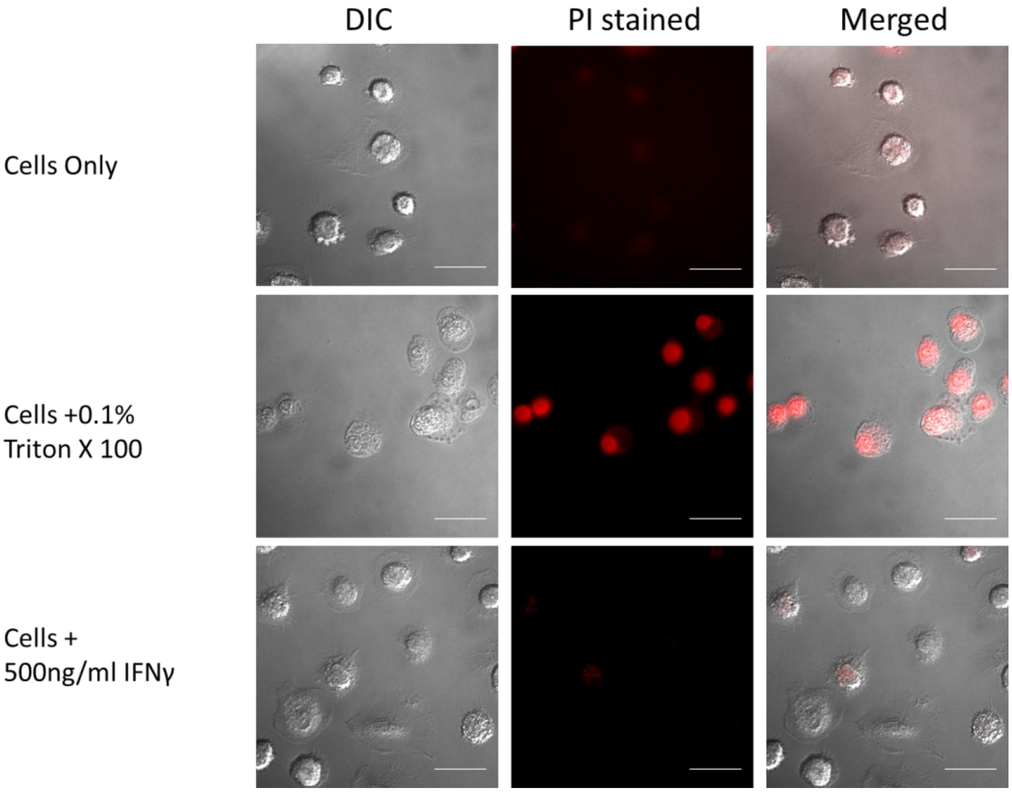


**Fig S5. High doses of IFNγ have minimal impact on macrophage viability.** Primary human macrophages were differentiated from blood. To ensure high dose IFNγ had little impact on cell viability cells were stained with propidium iodine (PI) to check for cell death. Differential interference contrast (DIC) microscopy was used to observe general cell morphology with red fluorescence used to determine the level of PI staining. Cells treated with IFNγ were observed after 24 h incubation and compared with cell in media only and a control positive for cell death, that had been treated with 0.1% triton X 100 10mins prior to microscopy.

**Table S1.** Bacteria and plasmids used in this study

| Strain/plasmid | Description | Reference/Supplier |
|---|---|---|
| *S. aureus* | | |
| COL | MRSA strain isolated in the 1960s (Clonal complex 8) | Gill et al. (2005) |
| COL Δ*seb* | Null mutant of *seb* in COL background | Xu et al. (2015) |
| COL Δ*seb* + pCM29::*seb* | COL Δ*seb* containing complementation plasmid expressing SEB | This Study |
| MW2 | MRSA strain isolated in 1998 (Clonal complex 1) | Fey et al. 2003 |
| MW2 Δ*sec* | Null mutant of *sec* in MW2 background | This Study |
| MW2 Δ*sec* + pCM29::*sec* | MW2 Δ*sec* containing complementation plasmid expressing SEC | This Study |
| *E. coli* | | |
| XL1-Blue | General cloning strain | Stratagene |
| SA30B | DNA methylation strain | Monk et al. (2015) |
| BL21 (DE3) | Protein expression strain | New England Biolabs |
| Plasmids | | |
| pKOR1 | Temperature-sensitive integration vector with inducible counter selection; Cmr | Bae et al. 2006 |
| pKOR1::*sec*_del | pKOR1 with *sec* flanking regions inserted, Cmr | This study |
| pCM29::*pLukM*-sGFP | pCM29 vector containing the active promoter of the leukocidin LukMF' | Vrieling et al. (2015) |
| pCM29::*pLukM-seb* | pCM29 vector containing the seb gene | This study |
| pCM29::*pLukM-sec* | pCM29 vector containing the sec gene | This study |
| pET28a | Protein expression plasmid, Kmr | Invitrogen |
| pET28a::*seb* | SEB protein expression plasmid, Kmr | Chau et al. (2009) |
| pET28q::sebN23A | $SEB_{N23A}$ protein expression plasmid, Kmr | Hayworth et al. (2012) |

[1] Cmr - Chloramphenicol resistant
[2] Kmr – Kanamycin resistant

**Table S2: Primers used in this study**

| Primer name | Sequence 5'-3' |
|---|---|
| SEB complementation | |
| pCM29-seb-kpnI-For [1] | CACAGGTACCAAAGGAGATAAAAAATGTATAAGAGATTA |
| pCM29-seb-EcoRI-Rev [1] | CACAGAATTCTCACTTTTTCTTTGTCGTAAG |
| SEC deletion[*] | |
| pKOR-sec-upstream-For [2] | **GGGGACAAGTTTGTACAAAAAAGCAGGCT**AGGCACAGCAATGTGTTCA |
| pKOR-sec-upstream-Rev[3] | GCTAGCACGCGTCTCCTTCATCCAACATTCCC |
| pKOR-sec-downstream-For[3] | ACGCGTGCTAGCGAGTGAAGATAGAAGTCCACCTTACA |
| pKOR-sec-downstream-Rev [2] | **GGGGACCACTTTGTACAAGAAAGCTGGGT**GCAAGCATCAAACAGTTACAAC |
| SEC-screen-For | GAAATCCTCTGTTTCTCCTTGAG |
| SEC-screen-Rev | CTATAAATATGGTTCTAACTCTC |
| SEC complementation | |
| pCM29-sec-kpnI-For [1] | CCGGTACCGTGTATCTAGATACTTTTTGGGAA |
| pCM29-sec-EcoRI-Rev [1] | CCGAATTCTTATCCATTCTTTGTTGTAAGGTGG |

[*] Primers design was based on primers previously described (Salgado-Pabón et al. 2013)

[1]Restriction sites (indicated in the primer name) are underlined in the primer sequence.

[2] attB sites used for recognition by BP Clonase are shown in boldface

[3] Overhang for splice PCR for fragment indicated are underlined in the primer sequence
